# Megafauna show pervasive yet distinct affinity to ocean fronts: the urgent need for adaptive conservation in a warming world

**DOI:** 10.1101/2025.06.17.660201

**Authors:** Isaac Brito-Morales, Boris Dewitte, Floriane Sudre, Christoph A. Rohner, Elliott L. Hazen, Kylie L. Scales, Matthieu Le Corre, Audrey Jaeger, Sophie Laran, Olivier Bousquet, Ana M. M. Sequeira, Tammy E. Davies, Daniel C. Dunn, Ronel Nel, Lee Hannah, Vincent Rossi

## Abstract

Fronts are ephemeral structures in the ocean that mark the boundaries between water masses of different properties, attracting a wide range of marine organisms, from plankton to whales. Despite their fundamental role in marine ecosystem functioning, the association with biodiversity has mainly focused on single species in regions with high data availability. Here, using multidecadal datasets on dynamical and thermal fronts, satellite tracking, and aerial observations, we assess marine megafauna associations with ocean fronts in the ecologically rich yet highly turbulent Mozambique Channel. We find that a diverse array of species associate with various ocean fronts, although the strength and type of affinity vary across taxa. Downscaled climate change simulations predict significant spatial shifts in front-rich areas by the end of the century. As climate change reshapes ocean front dynamics, adaptive management strategies will be essential to balance conservation and resource use in these critical ecosystems.

**Teaser:** Ocean fronts attract marine megafauna, but climate change might alter these habitats, requiring adaptive conservation strategies.

## Introduction

Ocean fronts are complex, three-dimensional dynamic features that separate water bodies of different properties, such as temperature and salinity. They form under various dynamical mechanisms and occur throughout the world’s oceans, varying over a wide range of temporal and spatial scales(*1*). Excluding the quasi-permanent fronts associated with topography, most current research focuses on mesoscale and submesoscale fronts in the upper ocean due to their impacts on climate change(*2*, *3*) and ecosystems(*4*). Ephemeral (sub-)mesoscale fronts can last from days to a few weeks and range horizontally from tens of meters to several hundred kilometers, with typical vertical extents of a few hundred meters(*5*, *6*). Some fronts feature strong vertical overturning circulations that modulate nutrient exchange between the nutrient-poor euphotic layer and the nutrient-rich deeper ocean, boosting primary productivity(*7*). Others tend to aggregate surrounding passively-drifting organisms, increasing planktonic food patchiness(*8*). Because ocean fronts act as biological hotspots in oligotrophic regions(*9*), they attract species of various trophic levels(*10*), including marine megafauna of conservation concern and commercially valuable fishes. This can lead to increased fishing activity in these areas(*11*, *12*) and conservation challenges, particularly through bycatch of non-target species and catch depredation by sharks and marine mammals(*13*, *14*). There is a pressing need for management strategies that balance fisheries sustainability with biodiversity conservation in these systems (*15*).

Here, we hypothesize that biophysical coupling at fronts (i.e., interaction between fronts and biodiversity) is primarily controlled by ephemeral niches created by the interplay between abiotic properties, such as temperature and currents, and the influence of these on species’ intrinsic behaviors, such as feeding, thermoregulation, navigation, and social behaviours. For instance, the Great Frigatebird (*Fregata minor*) navigates using Lagrangian coherent structures (i.e, LCS) to exploit the abundant food resources associated with biological aggregations at fronts(*16*).

Important commercial taxa like tuna have been observed using fronts to optimize thermal preference and foraging, i.e. exploiting the warmer sides of fronts to maintain higher body temperatures and then feeding more efficiently in the food-rich colder sides(*12*). Basking sharks (*Cetorhinus maximum*) navigate across different oceanic frontal structures, such as eddies and filaments, using isotherms as guides(*17*). Each of these species-specific utilizations are associated with different types of fronts (e.g., dynamical and thermal fronts), which can be assessed by different frontal metrics (e.g., Finite-Size Lyapunov Exponents [FSLE] and temperature horizontal gradients)(*18*).

Despite the affinity of marine megafauna to frontal systems, understanding the relationship between ocean-front typologies and biodiversity is still in its infancy, as is the conservation of frontal ecosystems. Previous research is dominated by single-species studies using unique front metrics and achieving low statistical significance(*19*). Studies of multiple species’ responses to multiple front dimensions integrated over large areas and long periods are rare, and almost entirely absent for the tropics (but see(*14*, *16*)). This is particularly pressing given that conflicts involving megafauna interactions with fisheries at ocean fronts are becoming an increasing conservation concern in tropical regions(*20*, *21*). At the same time, development of the Blue Economy seeks to improve income and food security by intensifying tropical fisheries(*22*).

Understanding multispecies interactions with fronts is crucial to identifying win-win scenarios between fisheries and conservation in the tropics that support ecosystem-based fisheries management. Aligning fishing efforts with the movement patterns of species that span various trophic levels at ocean fronts could reduce bycatch and other fishing impacts on marine megafauna(*23*) leading to ecosystem-wide benefits.

Understanding of ocean fronts requires an in-depth knowledge of the small-scale processes driving their formation and disintegration, as well as their long-term trends. This is essential for assessing their variability at meso- to submeso-scales, particularly as climate change is already altering ocean temperature(*24*), stratification(*25*), and surface currents(*26*), potentially affecting front frequency, intensity, duration, and associated biological responses(*27*, *28*). All of these aspects are critical for integrating ocean fronts into dynamic management frameworks. For example, understanding frontal variability can inform conservation planning and fisheries management by identifying how the change in ocean conditions might influence species distributions and ecosystem structure. Incorporating real-time ocean monitoring and predictive modeling could enhance management approaches, allowing for adaptive responses to shifting front dynamics(*29*).

Amongst the regions of the world where natural oceanic variability is the highest, the Mozambique Channel is embedded in one of the most energetic and turbulent Western boundary currents. Here, the intersection of the South Equatorial Current with the landmass of Madagascar, the scattered Islands and mainland Africa, drives the formation of mesoscale eddies and fronts that propagate southward through the channel towards the Agulhas Current(*30*). This channel is a remote and relatively pristine region of the ocean, currently being considered by several countries for a conservation planning initiative(*31*). Anthropogenic climate change affects the ocean in multiple ways, including warming and increased stratification(*32*), enhanced mesoscale variability(*33*), and reduced transport of the South and North East Madagascar Currents(*25*), potentially altering the spatial distribution, intensity, and frequency of fronts in the region. This, in turn, may impact frontal biological hotspots, affecting the abundance and distribution of predator species that rely on ocean fronts. A first step towards understanding ocean front habitats includes the assessment of: (i) front affinity by multiple species across trophic levels, (ii) front typology and its comparison through multiple frontal diagnostics, and (iii) the evolution of these biophysical relationships with climate change.

Here, we examine the use of ocean fronts by a wide array of marine megafauna (e.g., large elasmobranchs, seabirds(*34*), marine mammals, and sea turtles), using the Mozambique Channel as a study site. We test two distinct types of daily front diagnostics derived from remote-sensed archives: thermal and dynamical fronts. For thermal fronts, we utilize horizontal gradients computed with the Belkin & O’Reilly algorithm (BOA)(*35*) applied to high-resolution sea-surface temperature data. We capture dynamical fronts using Lagrangian Coherent Structures derived from backward Finite-Size Lyapunov Exponents (i.e., FSLE)(*33*, *36*). We combine data from satellite tracking and aerial surveys to document the interaction of marine megafauna with ocean fronts using these indices. We apply the front indices to multidecadal historical satellite records of daily front occurrences, examining the associations between multiple taxa and front intensity.

Assuming this association will persist, we then leverage the outputs of a nested downscaled regional climate simulation to project how the subregions prone to high frontal activity will evolve in the future. Our results provide unique insight into multispecies use of complex front structure in tropical waters, helping improve ecological understanding of front ecosystems and paving the way for better conservation management at fronts.

## Results

### Megafauna and ocean fronts: tracking data and aerial surveys

The spatio-temporal co-localization of ocean fronts with different taxonomic groups (i.e., seabirds, whale sharks, and turtles) reveal that marine megafauna associate closely with both dynamical and thermal fronts in the Mozambique Channel (Fig 1, 2). To analyze this association, we categorize the distance of these taxonomic groups to ocean fronts into 1km increments or ‘bins’, with the "0 bin" representing data points that fall exactly within a frontal area. In general, species from each taxonomic group are found in association with ocean fronts for both types of fronts. Seabirds associate with ocean fronts most closely based on both distance and frequency, followed by whale sharks and loggerhead turtles (Fig 1A, C, F, G). The significant aggregation of data in the 0 bin (i.e., a prominent peak higher than any other individual bin) suggests that a considerable proportion of the tracking data falls exactly within a frontal area for both front types. While whale sharks also exhibit this pattern for thermal fronts, the result is less pronounced for dynamical fronts (Fig 1A, E). The aerial survey data analyses corroborate the tracking data results, showing similar (although noisier) results (see Discussion). The corresponding density plots (Fig. 2) mirror these patterns.

**Fig. 1.**
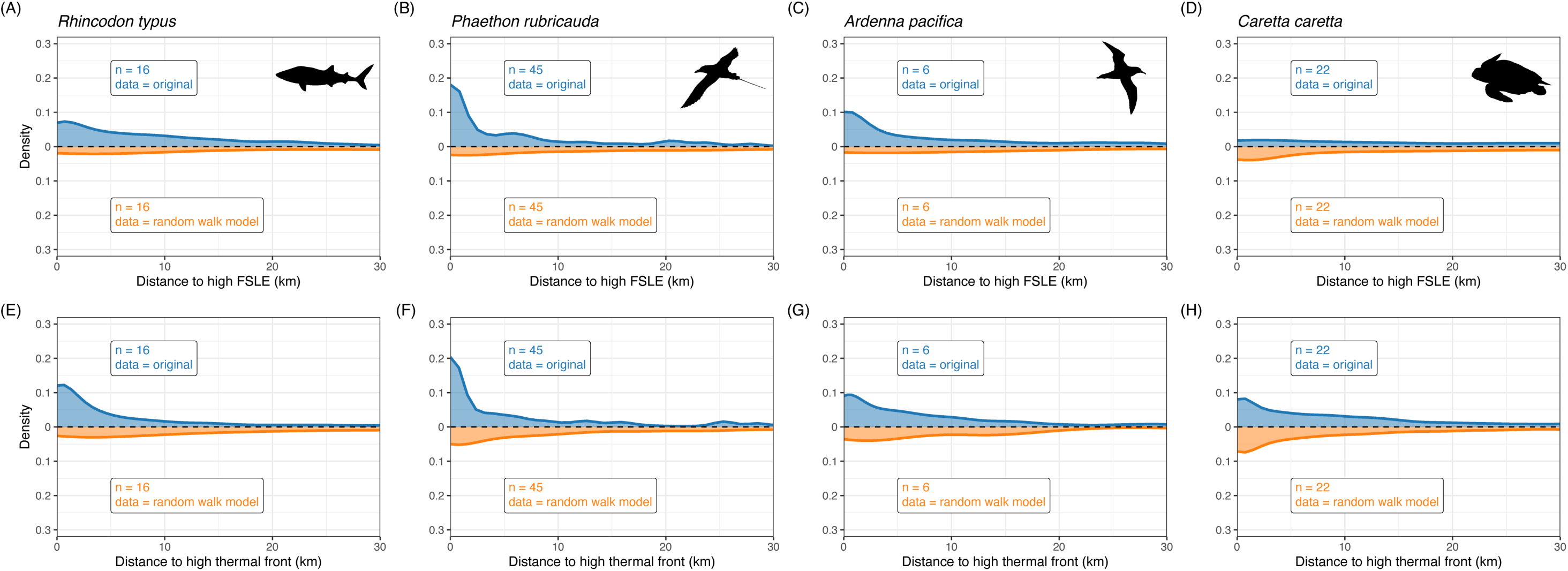
Kernel density plots illustrating the relationship between density and the distance to high-frequency front areas for three different taxonomic groups tracked with satellite-linked tags: (A, E) whale sharks (*Rhincodon typus*), (B, C, F, G) two seabird species: red-tailed tropicbird (*Phaeton rubicauda*) and wedge-tailed shearwater (*Ardenna pacifica*) , and (D, H) loggerhead turtle (*Caretta caretta*). In blue are the densities based on empirical tracks and in orange are densities based on randomised tracks. The upper row shows the distance (km) to high (upper quartile) Finite-Size Lyapunov Exponents (i.e. dynamical fronts). The lower row shows the distance (km) to high (upper quartile) thermal fronts derived from the Belkin and O’Reilly Algorithm(*35*).

**Fig 2.**
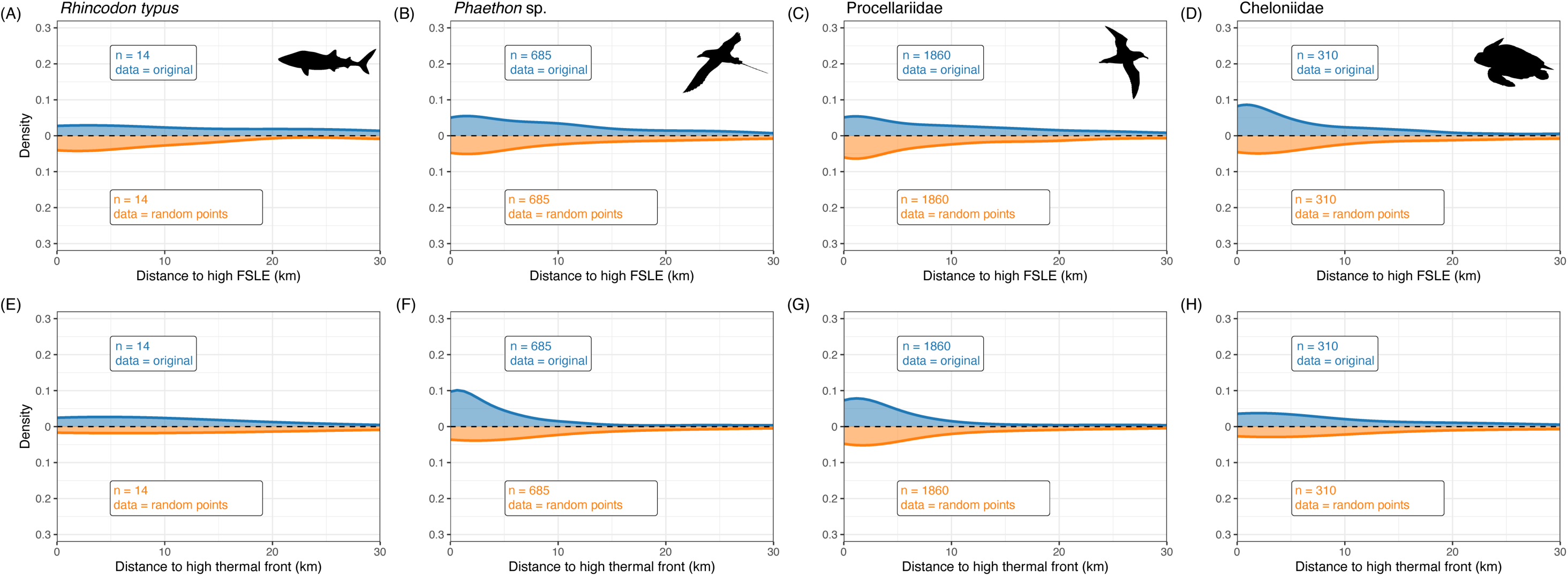
Kernel density plots illustrating the relationship between density and the distance to high-frequency front areas for three different taxonomic groups detected by aerial survey: (A, E) whale sharks, (B, C, F, G) two seabird species: tropicbirds undetermined (*Phaeton* sp.), and Procelladidae and (D, H) hard-shelled sea turtles. In blue are the densities based on empirical locations and in orange are densities based on randomised points. The upper row shows the distance (km) to high (upper quartile) Finite-Size Lyapunov Exponents (i.e. dynamical fronts). The lower row shows the distance (km) to high (upper quartile) thermal fronts derived from the Belkin and O’Reilly Algorithm(*35*).

### Random walks and random point analyses for megafauna

Since ocean fronts are common oceanographic features in the Mozambique Channel, the fact that biodiversity aligns well with them might be due to the high frontal frequency rather than a true selection process. To illustrate and evaluate this potential bias, we produced random walks for each tracking data set group (whale shark, seabirds, turtles) and random points along the survey track for aerial survey megafauna groups (cetaceans, seabirds, turtles, elasmobranchs) (see Methods section).

For the tracking dataset, our analysis reveals a consistent pattern in the random walk shapes across both dynamical and thermal fronts (Fig 1). Generally, for each megafauna group, the density for random walk does not exhibit a half-bell shape contrary to the observed data (Fig 1). This consistency in the random walk analysis strongly suggests that megafauna actively select ocean fronts. The observed tracks are not only different from the random tracks but also consistently closer to fronts, indicating that megafauna choose to move along fronts, with the exception of turtles along dynamical fronts (see Discussion section). The overlapping coefficient, which quantifies the similarity between distributions (where values near 0 indicate overlap due to chance and values near 1 indicate complete agreement), confirms this pattern. Values between the original and the random walk density distributions show a mean coefficient of 0.33 (Table 1), which indicates no overlap between the two distributions per each taxonomic group. This significant difference in the shape of the distributions implies that the two sets of data (i.e., original observations and random walk simulations) are fundamentally distinct.

**Table 1.** Similarity in the relationship between observed and randomly generated data by overlapping coefficients (see Methods) for both dynamical (i.e., FSLE) and thermal fronts, where near 0 represents overlap due to chance and near 1 represents complete agreement. For each species or group, we present the coefficients for tracking and serial datasets. Asterisks indicate the Kolmogorov-Smirnov (K-S) test significance levels in each comparison: (*) *p* < 0.05; (**) *p* < 0.01; (***) *p* < 0.001.

| Species / Group | Dynamical fronts (FSLE) | Thermal front |
| --- | --- | --- |
| **Tracking data** |  |  |
| Whale shark (*Rhincodon typus*) | 0.43*** | 0.36*** |
| Red-tailed tropicbird (*Phaethon rubricauda*) | 0.33*** | 0.35*** |
| Wedge-tailed shearwater (*Ardenna pacifica*) | 0.46*** | 0.47*** |
| Loggerhead turtle (*Caretta caretta*) | 0.45*** | 0.45*** |
| **Aerial observation data** |  |  |
| **Marine mammals** |  |  |
| Balaenopteridae | 0.53 | 0.10 |
| Sperm whale (*Physeter macrocephalus*) | 0.42 | 0.26 |
| Cuvier's beaked whale (*Ziphius cavirostris*) | 0.46 | 0.33 |
| *Mesoplodon* sp. | 0.29* | 0.46 |
| False killer whale (*Pseudorca crassidens*) | 0.27 | 0.36 |
| *Globicephala* / *Pseudorca* sp. | 0.36 | 0.44 |
| Short-finned pilot whale (*Globicephala macrorhynchus*) | 0.51 | 0.17 |
| Risso's dolphin (*Grampus griseus*) | 0.49 | 0.45* |
| *Peponocephala* / *Feresa* sp. | 0.25* | 0.32 |
| Melon-headed whale (*Peponocephala electra*) | 0.42 | 0.24 |
| Large delphininae | 0.45 | 0.35 |
| Bottlenose dolphin (*Tursiops truncatus*) | 0.37 | 0.35** |
| Small *delphininae* sp. | 0.39* | 0.29* |
| **Seabirds** |  |  |
| Noddies (*Anous* sp.) | 0.44 | 0.35*** |
| Frigate birds ( *Fregata* sp.) | 0.35 | 0.35* |
| Storm petrel (*Hydrobatidae* sp.) | 0.51 | 0.33 |
| “Grey” terns (*Onychoprion* sp.) | 0.40* | 0.37* |
| Red-tailed tropicbird (*Phaethon lepturus* | 0.47*** | 0.40* |
| Tropic birds (*Phaethon* sp.) | 0.43* | 0.32*** |
| *Procellariidae* sp. | 0.43*** | 0.41*** |
| Terns (*Sternidae* sp.) | 0.39*** | 0.37*** |
| Boobies (*Sula* sp.) | 0.38 | 0.39 |
| **Other Megafauna (turtles, rays, sharks)** |  |  |
| *Cheloniidae* sp. | 0.37 | 0.36 |
| *Dermochelys coriacea* | 0.50 | 0.47 |
| *Mobula* sp. | 0.24* | 0.43 |
| Rays (*Rajimorphii* sp.) | 0.36 | 0.27** |
| Whale shark (*Rhincodon typus*) | 0.12 | 0.23 |
| Sharks (Selachimorpha) | 0.34 | 0.45 |
| Hammerhead shark (*Sphyrna* sp.) | 0.33* | 0.40 |

For aerial surveys, the overlapping coefficient between the original shape and the random-point distribution shape shows a wide range of values across all taxa analyzed (Fig. 2, Table 1). In general, for both dynamical and thermal fronts, the simulated distribution exhibits a flatter or less pronounced peak compared to the observed distribution, suggesting poor agreement between the datasets. This pattern is generally consistent across each species within the different groups analyzed (cetaceans, seabirds, and other megafauna like turtles, rays, and sharks). This distinction can be better represented by comparing the patterns for the same taxonomic groups used in the tracking data analysis (Fig. 2).

Among cetaceans, the overlapping coefficient oscillates between 0.25 - 0.53 for dynamical fronts and 0.10 - 0.46 for thermal fronts. Only a few of the 15 taxa (Delphinidae and False killer whale, *Pseudorca crassidens*, for dynamical fronts, and Balaenopteridae for thermal fronts) show no significant association with fronts (Figures S1 and S2). Some dolphin taxa also show significant association with both front types; Risso’s dolphin (*Grampus griseus*) shows more affinity with thermal than with dynamical fronts (while both distributions are significantly different from the random ones), as does *Tursiops truncatus* (but in this case, only significant for thermal fronts).

For seabirds, the coefficients range from 0.35 to 0.51 at dynamical fronts and 0.32 - 0.41 at thermal fronts, with only three taxa showing patterns close to the empirical dataset for dynamical fronts (Grey-backed terns—*Onychoprion* sp.—Procellariidae, Sternidae) and only two families for thermal fronts (Hydrobatidae, Procellariidae, Figures S3 and S4). For these taxa, six out of the nine groups analyzed are significantly associated with both dynamical and thermal fronts (Table 1).

For turtles and elasmobranchs, the coefficients oscillate between 0.12–0.50 for dynamical fronts and 0.23–0.47 for thermal fronts (see Figures S5 and S6). While the observed patterns are more variable across taxa, they consistently show a similar distinction between frontal and non-frontal systems. In both cases, the random-point distributions are flatter than the observed distributions. Notably, only some taxa within the elasmobranch group show significant differences between the observed and random data, and this significance is not consistent across both frontal diagnostics (Table 1).

Finally, it is worth noting that some taxa exhibit bi- or multi-modal observed distributions (see secondary “bumps” for *Phaethon rubricuda* Fig. 1B, *Pseudorca crassidens* Fig. S1E, *Grampus griseus* Fig. S1H & S2H, *Balaenopteridae* Fig. S2A, *Peponocephala/Feresa sp.* Fig. S2I, *Fregata sp.* Fig. S3B, *Sula sp.* Fig. S4G, *Dermochelys coriacea* Fig. S5A & S6A, *Mobula birostris* Fig. S7B).

### Frequency of ocean fronts

A significant feature of the Mozambique Channel is the prevalence of ocean fronts, which occur in more than 30% of the whole South-Western Indian Ocean (SWIO) surface (Fig 3, Fig. S7A, B). Both dynamical and thermal fronts in the SWIO show regional differences in magnitude and frequency (Fig 3). Dynamic fronts are prominent within a large majority of the Mozambique Channel, with stronger magnitudes along the Mozambique shelf and south of Madagascar, as well as in the wake of the South-East Madagascar Current (SEMC) (Fig. S7A). Thermal fronts are frequent along continental shelf-breaks and within island wakes. They are notably occurring in the pathways of both the North-East Madagascar Current (NEMC) and SEMC, with overall more intense thermal fronts southward of 22°S (Fig. S7B). These findings arise from the interplay between intermittent physical oceanographic processes and established topography of the Channel.

**Fig. 3.**
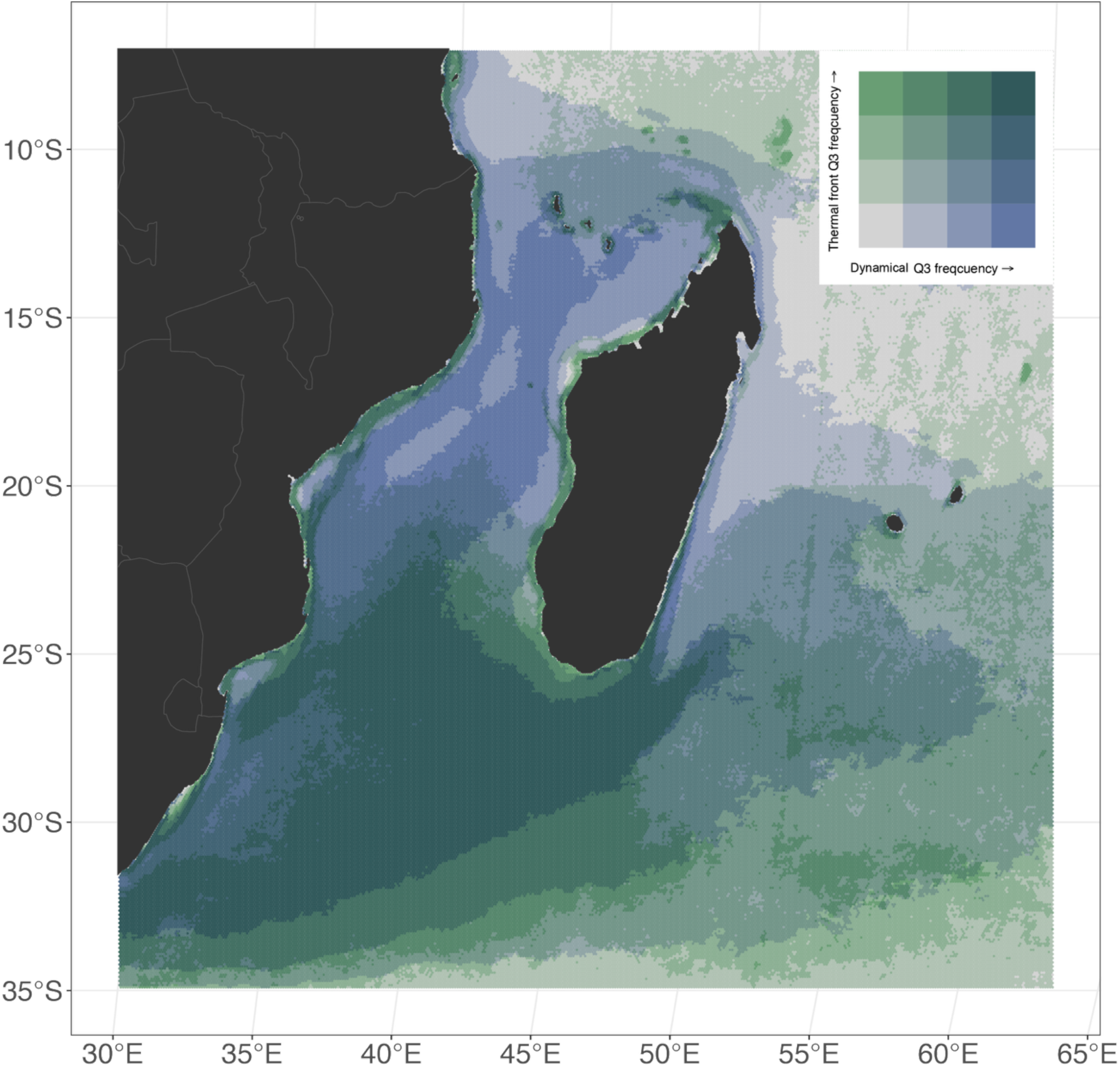
Spatial overlap of high-frequency occurrences (upper quartile) for backward-in-time Finite-Size Lyapunov Exponents (ridges of the FSLE field capture dynamical fronts) and thermal radients (ridges of the gradient field provide thermal fronts), across the South-Western Indian cean for the period 2003-2021, from NOMAD(*53*).

Overlapping the highly-frequent fronts of both typologies, reveals dynamical areas in which oceanographic processes could be especially relevant for marine megafauna and conservation (Fig 3). For instance, 15% of the SWIO experiences a high overlap of high-frequency dynamical and thermal fronts. These areas are mainly located along the coast of Mozambique, along the northeast coast of Madagascar, throughout the central and southern regions of the Mozambique Channel, and also near the formation region of the Agulhas Current (Fig 3). However, the spatial distribution of both front diagnostics does not overlap homogeneously: while 4.2% of the area shows high dynamical front characteristics, these areas simultaneously exhibit low thermal front intensity. This suggests a complex interplay between different types of ocean fronts in the SWIO. It also demonstrates that, while both front diagnostics highlight consistently some front-rich regions (e.g. island wakes, pathways of NEMC/SEMC, as well as the southern MC as a whole), they also exhibit different abilities in capturing front habitats. For instance, shelf frontal activity is better captured by thermal fronts while southward travelling frontal eddies in the MC appear more robustly depicted by dynamical fronts.

### Ocean fronts and climate change

Areas where tight associations between fronts and marine megafauna occur may shift with climate change, as warming, circulation changes and increased stratification could affect the location, magnitude and persistence of both thermal and dynamical fronts. To illustrate the associated complexity, we performed a regional dynamical downscaling of climate projections of a global Earth System Model under a high-emissions scenario (see Methods for more details) to estimate mean changes in frontal dynamics under future climate following (*37*). We have focused our analyses on two main aspects: changes in thermal and dynamical front frequency between past (1993–2014) and future decades (2076–2097), as well as changes in ocean-front frequency within regions exhibiting high occurrence rates in past decades (Fig 4). Note that the warming magnitude and patterns simulated by our regional model (Fig. 8) are consistent with those reported by (*32*); moreover, changes in surface current, such as the slowdown of the Mozambique Channel, Northen Mozambique Channel and Southern Mozambique Channel are coherent with (*26*).

**Fig. 4.**
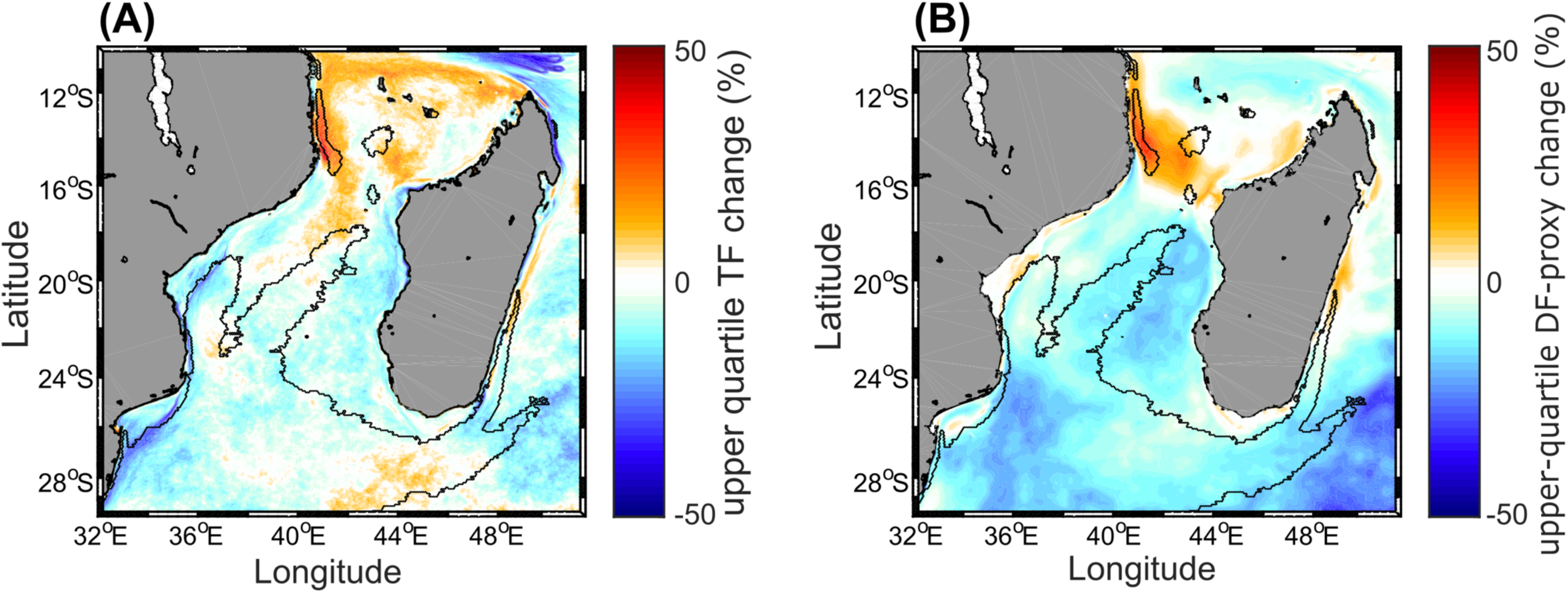
Changes in upper-quartile thermal front (left) and upper-quartile dynamical front (right, through a proxy, see Methods) frequency (in % of the upper-quartile front frequency in the hindcast) between hindcast and pseudo global warming experiments. The bold black contour delineates the Areas of High Front Frequency (AHFF) derived from both front diagnostics in the NOMAD, also corresponding to the darker areas in Fig. 3.

Our results reveal clear spatial contrasts in the projected changes of both thermal and dynamical front frequencies (Fig. 4). In general, increases in upper-quartile front frequencies are observed in parts of the northern Mozambique Channel (north of ∼17°S) and southwest of Madagascar— especially for thermal fronts—while decreases are more prominent in the southern channel and around the SEMC retroflection, particularly for dynamical fronts. This meridional divide aligns with previously observed trends of mesoscale activity under climate change (*38*, *39*), although the spatial extent of these zones varies slightly between front types. Notably, areas showing substantial climate-driven shifts in both thermal and dynamical fronts overlap with known megafauna habitats reported in earlier studies(*40*, *41*).

Finally, when focusing on the area characterized by high frontal activity in past decades (covering approximately 1,122,711 km²; black contours in Fig. 4), our climate change model experiment shows that this region is projected to experience significant changes in both thermal and dynamical front occurrence. For instance, thermal fronts are projected to increase by around 5% in about 26% of this area—mainly in the Comoros Basin and south of Madagascar. However, the remaining 74% of the region is expected to experience a mean 7% decrease in thermal front activity, particularly in the central and southern Mozambique Channel (Fig. 4A). Similarly, for dynamical fronts, only about 5% of this historically front-rich region is projected to show a 10% increase—especially near the Comoros Basin and the eastern Madagascar shelves—while the remaining 95% will exhibit a mean 12% decrease, concentrated in the central and southern Mozambique Channel (Fig. 4B).

## Discussion

Our results demonstrate, for the first time, that multiple marine megafauna species—including seabirds, turtles, and whale sharks—actively select ocean fronts as key ecological habitats. Both thermal and dynamical fronts are important to multiple species, and potentially for multiple behavioral clues, as suggested by the multi-modal distributions of some species. Indeed, megafauna association with fronts is not uniform across taxa—seabirds exhibit the strongest affinity for frontal systems, followed by cetaceans, while turtles show the most variable response, highlighting species-specific interactions with ocean dynamics. These findings build on previous studies that have demonstrated the ecological significance of ocean fronts as foraging zones for top predators, such as Great Frigatebirds(*16*), and as areas of high fishing activity that elevate bycatch risks for sharks, sea turtles, and cetaceans(*14*). In the California Current, similar features have been linked to fisheries interactions, reinforcing the need for dynamic conservation measures(*42*). Moreover, our findings reveal that 15% of the SWIO experiences a high overlap of both thermal and dynamical fronts, suggesting that these dynamic oceanographic features create important habitats for megafauna. Altogether, it demonstrates that a wide variety of species depend on these ephemeral ecosystems, underscoring the ecological importance of these regions here and elsewhere for conservation.

Our study highlights how species’ life history can influence the observed patterns of how megafauna interact with fronts, underscoring the importance of considering ontogenetic and behavioral components when interpreting this association. For example, tracking data on turtles shows no apparent interaction with fronts (Fig. 2D, H). Although this contrasts with previous studies(*43*, *44*), the lack of a pattern could be attributed to the life stages (e.g., young adults) of the individuals tracked for their first breading events(*45*) or the fact that sea-turtle navigation use diverse cues such as chemical or geomagnetic(*46*). Other species, such as whale sharks show strong associations with ocean fronts here (Fig. 1A, E) and in previous studies(*47*), but aerial observations do not provide such clear evidence ( Fig. 2A, E, Fig. S5, S6). The same applies for other marine megafauna in aerial surveys — some species or groups show affinity for fronts, but the pattern is less pronounced compared to the tracking data (Fig. S5, S6). Here, it is important to note that, unlike tracking data, aerial observations are conducted along predetermined line transects and do not necessarily capture the same species/individual. Another key difference is that they record individuals only when they are at the surface. Thus, the observations for marine mammals, turtles, and elasmobranchs (not necessarily birds) (Fig S1, S2) could be biased due to individual-specific representation but most importantly to surface-only observations. This highlights the importance of combining different data sources (i.e., tracking, aerial observations, systematic surveys) in data-poor regions to improve our appraisal of megafauna/front interactions while caution is needed to consider the limitations and biases inherent in each dataset.

We have provided evidence that not only are species using ocean fronts, but also that species use different types of ocean fronts. Both diagnostics reveal different aspects of the ocean dynamics, as thermal fronts are based on the Eulerian analysis of an active oceanic tracer field while dynamical fronts originate from the Lagrangian perspective of passive tracers, which makes the analyses complementary to each other. It is also important to note that both diagnostics used here have distinct biases. For example, while thermal gradients highlight semi-permanent fronts near continental shelf-breaks and other bathymetric features, these may be missed by dynamical fronts as due to the artificial grounding of particles onto surrounding shorelines. Conversely, cloud cover, especially during the austral winter(*39*), generates gaps in remote-sensed sea surface temperature products, negatively affecting the computations of thermal fronts but not of dynamical fronts, which are themselves derived from currents combining altimetry and scatterometry. In addition, chlorophyll-a fronts, which reflect surface productivity gradients, represent another type of ocean front that has been linked to marine megafauna, particularly in areas where primary productivity drives prey aggregation(*48*). This intricacy of oceanic physical and biogeochemical processes calls for further research in ocean front detection, which, as this study and previous ones suggest, creates productive habitats that attract diverse marine megafauna (*15*, *16*, *48*).

Our climate change simulations show significant spatial shifts in front-rich areas by the end of the century. The decrease in both dynamical and thermal fronts activity, mainly in the Southern Mozambique Channel, indicates that these key areas for megafauna might become less suitable in the future. This could result in megafauna shifting their distribution patterns, which might have implications in current conservation management strategies. Conversely, areas such as the northern Mozambique Channel, where frontal activity is projected to increase, might become even more important for biodiversity conservation. The strong overlap between these areas and megafauna occupancy reinforces the need for adaptive management strategies that consider not only current megafauna distribution estimates(*38*), but also future shifts of their preferred habitats driven by climate change. Incorporating climate projections into MPA planning and area-based management(*49*) will be essential for ensuring the long-term effectiveness of conservation efforts(*27*, *50*).

There are several caveats in our study that should be considered. First, our study uses ecological metrics based solely on physical aspects of ocean circulation to demonstrate that ocean fronts aggregate marine megafauna(*51*). However, it is possible that megafauna aggregation at fronts may also depend on several other factors not considered here, such as oxygen levels, acidity, and abundances of actively-swimming preys (*5*, *6*, *27*). Second, our study does not attempt to identify specific behavioral patterns for the evaluated taxonomic groups. Note however that our methodology could allow investigating such effects, as suggested by the multi-modal distributions indicating that, behind the general affinity for fronts, distinct ecological behaviors may occur at various distances from the nearest fronts. While we acknowledge that species might use fronts for several purposes(*52*), our focus is on highlighting the significance of these dynamic features for marine biodiversity, rather than explaining the underlying behavioral patterns. Third, our definition of an ocean front is based on specific criteria: we use backward Finite-Size Lyapunov Exponents (FSLE) and thermal gradient values that fall in the upper quartile of their respective distributions (refer to the Methods section for details). This generic threshold approach performs well in highlighting dynamical and thermal fronts relevant to megafauna, yet it depends on the domain and time period considered. Other useful approaches involve area-specific thresholds based on ecological regionalization(*53*). Fourth, our study relies on surface-only fields (seawater temperature and horizontal currents) to estimate dynamical and thermal fronts. A significant limitation here is that fronts also have a subsurface component(*53*), which is not captured in our study, potentially affecting the comprehensiveness of our findings. For instance, air-breathers such as seabirds and cetaceans may use different parts of the water column around fronts compared to deep-diving species like tuna, billfish, or sharks. This might explain why surface- orientated taxa, such as seabirds, had a stronger association with fronts than other taxa that frequently dive or forage at depth (e.g. whale shark, sperm whale, Physeter macrocephalus).

Future studies could address this limitation by incorporating data-assimilative ocean models, which provide more accurate approximations of subsurface front structures.

Deducing megafauna affinity for frontal systems from the approach we developed here opens promising prospects for marine conservation. Future research should focus on improving predictive tools that incorporate ocean front dynamics, allowing for their formal integration into marine spatial planning. Better accounting for these dynamic features in marine conservation would enhance the effectiveness of traditionally static approaches, leading to more adaptive and ecologically relevant conservation frameworks(*54*, *55*). Finally, ocean fronts involve several dynamical and thermo-dynamical processes in their formation and dissipation, leading to diverse diagnostics for detecting these structures. While our study is pioneering in considering frontal events from both Eulerian and Lagrangian perspectives and comparing them with multispecies megafauna data, future research could benefit from including fronts of chlorophyll-a concentration or primary productivity(*56*), to better link physical diagnostics with food web spatial dynamics. Another still controversial question is whether organisms’ affinity to fronts operate simultaneously at all trophic levels (albeit various passive and/or active mechanisms), potentially indicating that fronts boost in-situ trophic interactions(*10*). While recent research(*52*) focusing on mid-trophic level organisms challenges the previously-suggested oasis effects of fronts, probably due to the use of an outdated detection method, our study demonstrates that conclusions may differ depending on the type of frontal diagnostics.

Since ocean fronts shape marine biodiversity in subtropical marine ecosystems(*57*), adaptive protection measures for frontal zones could help expand the current static ocean conservation paradigm(*58*). Dynamic marine conservation at intra- and inter-annual time-scales has recently emerged as a potential solution for highly mobile biodiversity(*54*, *59*); these results dealing with decadal time-scales could have positive implications for it. For example, one potential dynamic marine protected area (MPA) approach should be to establish a well-connected network of MPAs that accounts for the spatial variability of front-rich regions and their decadal changes in the Mozambique Channel . By strategically placing MPAs in areas where the most pronounced ocean fronts occur more frequently, we can help protect the critical habitats that support the diverse marine life in the region. Another approach would be to use ocean-front data to inform MPA design and management, ensuring MPAs encompass areas of high ocean-frontal frequency, with management strategies that adapt to changing dynamics over decades. French authorities have already gathered frontal statistics(*31*), alongside other biological and topographical datasets, and analyzed expert interviews to prioritize factors and delineate MPA boundaries as objectively as possible, given current knowledge. However, this process did not account for future changes yet. Incorporating knowledge of ocean fronts and of their future evolution into MPA design and management could ultimately help ensure the long-term sustainability of marine ecosystems in the Mozambique Channel.

This study highlights the ecological importance of ocean fronts and the need to balance conservation efforts with sustainable human activities. While we have shown that megafauna use both thermal and dynamical fronts, further research is necessary to explore the potential for spatial and vertical partitioning between fisheries target species and non-target megafauna.

Understanding these patterns could inform dynamic management approaches and fishing strategies that minimize conflicts and enhance both conservation and fisheries productivity. Dynamic management approaches that can predict megafauna and target species habitat in space and time have been shown to allow increased catch rates while meeting conservation objectives(*60*). Although this specific aspect is beyond the scope of our current study, it aligns with the pressing need to adapt resource management in the face of climate change. This work lays a foundation for future research and policy aimed at protecting these dynamic ecosystems, ensuring both their ecological integrity and their continued contribution to human well-being.

## Materials and Methods

### Study region

Our study focuses on the Mozambique Channel as a data-poor case study for managing tropical ocean fronts for change, now and in the future. The Mozambique Channel, situated in the South- Western Indian Ocean (SWIO) between Madagascar and the African continent, is a region known for its rich ephemeral ocean fronts(*51*, *53*). This area exhibits a complex and variable surface and sub-surface circulation predominantly characterized by mesoscale activity(*38*, *61*). Successive trains of mesoscale eddies propagate southward from the entrance of the channel at 12°S towards the beginning of the Agulhas current around 27°S(*30*). In the Channel, ephemeral fronts form in-between those eddies and on the edge of the Northeast and Southeast Madagascar Currents(*51*). The Channel has also been described as a superhighway for many marine species(*62*), which use it as corridors during specific times of the year depending on the species considered. These species are known to rely on the Channel as vital habitats for activities such as feeding, mating, and socializing, particularly during migration periods(*62*).

### Dynamical and thermal fronts

Daily maps of ocean fronts are derived from 2-dimensional surface fields of oceanic variables. They are usually defined in a Eulerian framework by highlighting, at each time increment, horizontal gradients in either density, temperature, salinity, chlorophyll-a or sea-surface height. Eulerian fronts have been widely used in marine ecology and fisheries to explain various biological spatial patterns(*63*). Focusing on temperature, surface thermal fronts are captured here by looking for high values in thermal gradient fields computed from Sea Surface Temperature (SST).

Ocean fronts have been also characterized by peculiar dynamical signatures, such as flow convergence and along-front jets, which can be detected in a Lagrangian framework from 2D horizontal flow fields(*64*). Here we use Attracting Lagrangian Coherent Structures (LCS), which are revealed by ridges of backward-in-time Finite-Size Lyapunov Exponents (FSLE) and form transport barriers along which passive tracers tend to converge(*33*, *36*). As such, they highlight accumulation places of prey, including planktonic organisms and other poorly mobile micronekton species, subsequently attracting small and large pelagic organisms and other apex predators. Indeed, they have been already shown to influence the structure and functioning of marine ecosystems at various trophic levels(*8*, *14*, *16*, *65–67*).

In this work we combine both Eulerian and Lagrangian perspectives, which we refer to as thermal and dynamical fronts respectively, to offer a comprehensive diagnostic of ocean fronts and how they relate to marine life. More specifically, ocean fronts for past decades (used to pair with animal’s observations) have been derived from the oceaN frOnt dataset for the Mediterranean seA and southwest inDian ocean(*53*). It provides high-resolution daily thermal gradients and backward FSLE fields for the SWIO between 2003 and 2021 computed from satellite data.

Thermal gradients are expressed on a 1/100° grid (approx. 1 km) in °C km^-1^, while backward FSLE values are given on a 1/64° (approx. 1.56 km) grid in day^-1^. Thermal gradients are computed from the Multi-scale Ultra-high Resolution SST analysis(*68*) (accessed on Jan 10th, 2023) with the Belkin and O-Reilly Algorithm(*35*). Backward FSLE are computed with (*33*)’s algorithm from satellite-derived surface currents, which combine Altimetric absolute geostrophic velocities and Modeled Ekman Current Reprocessing at 0.25° spatial resolution. Ocean fronts for future decades as well as for the change analyses have been computed on modelled surface total currents and temperatures generated by regional state-of-the-art hydrodynamical models (see next subsection). We define thermal and dynamical fronts as the upper quartile values of the daily magnitude of thermal gradient and backward FSLE distributions (respectively). Then for both front diagnostics, the upper-quartile frequency is computed as a percentage of days in which each pixel is found in an upper-quartile front state.

### Megafauna data: satellite tracking

Seabirds: GPS loggers were used to track the at-sea movements of two seabird species, the red- tailed tropicbird (*Phaethon rubricauda*) breeding on Nosy Ve (23.65°S, 43.60°E, Madagascar), and the wedge-tailed shearwater (*Ardenna pacifica*), breeding on Reunion Island (21.37°S, 55.55°E, France). The loggers were attached to tail feathers of incubating birds using TESA® tape (Beiersdorf, AG, Germany) and were retrieved between 3 and 21 days after deployment.

Data on the movements of 45 red-tailed tropicbirds were collected from May 2018 to March 2019 and data from 6 wedge-tailed shearwaters were gathered from November 2017 to January 2019. Three types of GPS loggers were used: i-gotU GT-120 (15g, Mobile Action Technology Inc), CatLog Gen1 (20g, Perthold Engineering LLC) and CatLog Gen2 (18g, Perthold Engineering LLC). Tags were set to record a GPS location every 10 minutes. Bird handling and tag deployments were performed as part of a scientifically and ethically approved research program (*41*).

Sharks: Whale sharks are classified as endangered species on the IUCN Red List, with fishing, bycatch and ship strike as the main contemporary threats. Whale shark movements were tracked at the two main aggregation sites for the species in the Mozambique Channel, off Praia do Tofo, southern Mozambique, and Nosy Be, northwestern Madagascar. Fifteen juvenile whale sharks (12 males, 3 females, size range = 540–865 cm total length [TL]) were tracked from Praia do Tofo in 2010-12(*69*) and eight juveniles (6 males, 2 females, 400–700 cm TL) were tracked from Nosy Be in 2016/17(*70*). We attached SPOT-5 satellite tags (Wildlife Computers) to whale sharks on a ∼1.8 m long tether by placing a titanium dart in the thick skin near the 1st dorsal fin while snorkeling alongside free-swimming whale sharks. The positively-buoyant tag sends location transmissions via the ARGOS satellites whenever the shark is at the surface and the tag is exposed to air, and a satellite is overhead. In Mozambique, we only used locations from transmission classes 3, 2 and 1 which are the most accurate and have errors of ∼0.49, 0.94 and 1.1 km, respectively(*71*). In Madagascar, we used the Douglas filter to remove inaccurate locations but keep some locations from classes 0, B and A. We removed short tracks for this analysis, leaving 8 tracks that departed from Mozambique and 8 tracks from Madagascar.

Turtles: Argos and GPS satellite tags were relied upon to track the movements of 22 sub adult loggerhead (*Caretta caretta*) marine turtles equipped from Reunion Island in the frame of an international research program (*72*) over the period 2019-2021. A Kalman filter scheme was used to process Argos data after discarding classes B and Z locations and a maximum travel rate of 10 ms-1 was applied to filter larger outliers arising from the lognormal distribution of positioning errors. Animal positions were further refined using the “FoieGras” library, which was set up to resample animal trajectories at regular time steps of 6 to 12-hours(*72*).

### Megafauna data: aerial survey

Aerial surveys were carried out from mid-December to the beginning of April 2010, following a standard line-transect methodology. A multi-taxa protocol was applied to collect data on marine mammals, turtles, elasmobranchs, and large fishes. Nonetheless, seabird sightings were collected following a strip-transect methodology (within a 200-meter strip; see(*40*) for more details on the protocol). Species identification was made to the lowest taxonomic level whenever possible, but groupings were inevitable for several taxa that could not be distinguished from the air. For marine mammals, 13 groups were considered: small Delphininae undetermined (*Stenella* sp.), large Delphininae, including bottlenose dolphin, small Globicephalinae undetermined pygmy killer whale *Feresa attenuata*, melon-headed whale (*Peponocephala electra*), Risso’s dolphin, large Globicephalinae undetermined (short-finned pilot whale - *Globicephala macrorhynchus*-, killer whale -*Orcinus orca*-, False killer whale -*Pseudorca crassidens*), beaked whales (*Mesoplodon* sp. or Ziphiidae), sperm whales and Balaenopteridae,. For seabirds, seven groups were comprised: brown terns, grey terns (*Onychoprion* sp.), noddies (*Anous* sp.), petrels and shearwaters (Procellariidae), tropicbirds (*Phaeton lepturus* and P. sp.), boobies (*Sula* sp.), and frigatebirds (*Fregata* sp.; cf. Supplementary Table 2 for the list of species). Finally, a “hard-shelled group” (Cheloniidae) and leatherback turtles (*Dermochelys coriacea*) were considered, in addition to manta rays, unidentified rays, whale sharks, hammerhead sharks, and unidentified sharks.

### Megafauna at ocean fronts

To assess whether megafauna closely follow ocean fronts, we calculated the distance (in kilometers) between each track location and aerial observation with the upper quartile values of the corresponding daily magnitude of thermal and dynamical fronts. If megafauna is actively present at these ocean fronts in the Mozambique Channel, we would expect most of the distances for each individual in both datasets to be zero.

### Random walk analysis for tracking data

To test whether the positioning of each individual from three marine megafauna groups (whale sharks, seabirds, and turtles) was biased towards the high frequency of ocean fronts (both dynamical and thermal fronts) in the Mozambique Channel, we used Correlated Random Walks (CRWs) for each set of tracking data. The initial position for each random walk was aligned with that of the corresponding focal individual. The duration of each random walk was tailored to the specific movement frequency of each individual. The Correlated Random Walk (CRW) can then be mathematically formulated as:

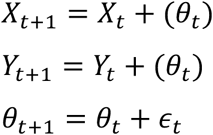

Here, *Xt* and *Yt* represent the coordinates of the individual at time t, and θt indicates the direction at time t, which is correlated with the direction at time *t-1*. The term εt is a random error introduced at each step to induce correlation, typically drawn from a normal distribution centred around zero. We use a random angle condition (*θ*) and a high correlation (ρ=0.9) value to each randomization. The high correlation ensures that the change in angle between steps is minimal, maintaining a relatively consistent direction from one step to the next. This approach allowed us to simulate realistic movement patterns of the studied species and assess their potential preference towards ocean fronts in the Mozambique Channel. All computation was undertaken using R 4.2.3(*73*).

### Random points for aerial survey data

As described above (Methods: Observational Data), the aerial survey data consisted of airborne observational dataset of several megafauna groups (see Table S1) alongside different transects in the Mozambique Channel and the Western Indian Ocean(*40*). To test the hypothesis if the observed aerial pattern was also biased towards ocean fronts (both dynamical and thermal fronts), we created random points for the study area. Random points were generated in alignment with the transect positions of the original observations for each taxon. This approach ensured that the random points were confined within the geographical limits of the study area, mirroring the locations where actual organisms were observed. This method helped to account for the spatial-temporal bias of the heterogeneous sampling of the frontal fields by the plane transects. The generation and analysis of these data points, as well as all other computations, were conducted using R version 4.2.3(*73*). Detailed information about the dataset and methodology can be found in(*40*).

### Climate change analysis

We evaluate the impact of climate change on the spatial patterns of front-rich surface habitats in the Mozambique Channel based on decadal experiment with a regional oceanic model under different forcing conditions. The regional model is the Coastal and Regional Ocean COmmunity model (CROCO(*74*)) built upon the Regional Ocean Modeling System (ROMS)(*75*). The model configuration consists in two embedded domains in a one-way (offline) nesting setup: a parent domain at a 1/12° horizontal resolution consisting in a rectangular basin over the region (27.1°E– 54.2°E, 7.5°S–33.1°S), and an inner child domain at higher-resolution (1/36°) over the region (30.6°E–51.9°E, 9.8°S–29.7°S). Sixty sigma vertical layers are used that are stretched vertically for increased resolution at the surface boundary layer. We use the GEBCO 2014 bathymetry(*76*) for consistency with the background lateral boundary conditions that are derived from the outputs of a ROMS simulation at 1/4°. We utilize two 22-year high-resolution (e.g. 1/36°) regional simulations (hereafter “control” and “projection”), each differing in their boundary conditions, which correspond to either contemporary or future climate scenarios. A 3-year spin-up period is included (but not analyzed) to allow model stabilization.

The baseline, or “control” run, consists in running the regional child model over the period 1993- 2014 using boundary forcings of the contemporary climate derived from the hindcast experiment detailed by Sudre et al.(*51*). Bulk formulations are used to derive atmospheric forcings from the model oceanic currents and SST and the ERA (ECMWF Re-Analysis) interim daily-mean fields (*77*). This simulation is comparable to that analyzed in Sudre et al.(*51*) , where simulated currents and temperatures have been extensively validated against remote-sensed and climatological observations. It also accounts realistically for the mean patterns of eddy activity, as shown by the analogous spatial patterns (spatial correlation coefficient reaches 0.76, Fig. S9A, B). Note that the regional model simulates slightly larger amplitude of mean Eddy Kinetic Energy (EKE) than satellite-derived EKE, potentially due to model biases or the coarser resolution (1/4°) of altimetric data that tends to underestimate in-situ EKE.

In a so-called pseudo global warming experiment, or “projection”, we add a climate warming perturbation (i.e., delta) to both atmospheric and boundary forcings of the control run(*78*). The delta consists in the difference of the climatology of each variable in the boundary forcing between the future and present climates as simulated by the Community Earth System Model Large Ensemble (CESM-LENS)(*79*). This model resource has been widely used for climate studies due to the skill of the model in accounting for many aspects of the Indo-Pacific tropical climate variability(*80*) and the availability of numerous realizations allowing for a robust estimate of anthropologically forced changes.

The mean conditions of the present climate are approximated by the climatology over 30 years of the historical scenario while the future climate corresponds to the Representative Concentration Pathway (RCP) 8.5 (high emission scenario for "business as usual”) scenario for the period 2076- 2097. The delta thus accounts for the change in seasonality of the main variables between the future and present climates, which is incorporated in the regional model for the sensitivity experiment. This approach is commonly used in the regional modeling community to deal with biases in global climate models(*81*). Forty members of the CESM-LENS are used to estimate the climatologies for each mean climate conditions and thus the delta, which ensures that the effect of natural variability in the boundary conditions is marginal in this experiment. This protocol enables the simulation of a climate change pattern consistent with that of the global model (See Supplementary Material), while incorporating enhanced physical realism through the inclusion of mesoscale dynamics. Indeed, while the downscaled regional model simulates a slightly cooler overall SST changes than global models (that could be attributed to enhanced mixing from more intense mesoscale activity), it does capture much finer spatial pattern changes (Fig. S9C, D).

Front changes were evaluated in both baseline and pseudo global warming model experiments on the 1/36° grid, at surface. We compute daily thermal gradients with the Belkin and O’Reilly Algorithm(*35*) and daily intra-seasonal Eddy Kinetic Energy (EKE) following the method described in (*33*), with:

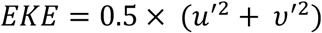

Where the daily turbulent eastward and northward velocities u’ and v’ (respectively) are defined as deviations from the temporal mean of total currents.

EKE serves as a proxy of the dynamical fronts derived from ridges of the backward-in-time FSLE field based on a regional statistical empirical relationship published previously by Hernandez Carrasco et al.(*33*) (their Fig. 8h). In the MC, backward FSLE and EKE are interrelated following:

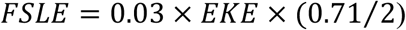

For both front diagnostics, the upper-quartile frequency is considered by computing the percentage of days in which each pixel is in an upper-quartile front state. The upper quartile front state is defined as the top 25% of the distribution (in either thermal or dynamical fronts) over the entire domain and period of both present and future simulation.

## Acknowledgments

We thank Simon Pierce, Stella Diamant, and Clare Prebble for their help with tagging whale sharks. Field assistance for seabird tracking data collection was provided by Anasvaler Mbelomanana, Jade Lopez, and Vincent Hoarau.

## Funding

National Science Foundation grant NSF-2029710 (I.B.M., L.H.)

French National Research Agency grant ANR-20-BFOC-0006-04 (F.S., V.R.) European FEDER Fund grant 1166-39417 (V.R.)

Excellence Initiative of AMU - AMIDEX ("Investissements d’Avenir") (V.R.) European Space Agency grant 4000141547/23/IDT (V.R.)

OMNCG Federation of the University of Réunion (SPY program) grant (M.L.C., A.J.) Waterlust, Aqua-Firma, and the Shark Foundation grant (C.R.)

Australian Research Council Discovery Early Career Researcher Award DECRA 2021 DE210100367 (K.S.)

Agencia Nacional de Investigación y Desarrollo (ANID) grant R20F0008-CEAZA (B.D.) Agencia Nacional de Investigación y Desarrollo (ANID) grant FB210021 (COPAS COASTAL) (B.D.)

Australian Research Council Discovery Project grant DP210103091 (A.M.M.S.)

## Author contributions

Conceptualization: I.B.M., L.H., V.R. Methodology: I.B.M., F.S., B.D., V.R. Writing—original draft: I.B.M.

Writing—review & editing: All the authors contributed equally to discussion of ideas and commented on the manuscript.

## Competing interests

Authors declare that they have no competing interests.

## Data and materials availability

The original satellite tracking dataset used in this study (seabirds, whales, sharks, and turtles) is available upon request. However, a curated version has been included in the GitHub repository: https://github.com/IsaakBM/mmw_belmont. Aerial survey datasets are freely available online at https://pelabox.univ-lr.fr/pelagis/PelaObs/. All code used in this study is also available in the GitHub repository: https://github.com/IsaakBM/mmw_belmont. Satellite-derived front datasets (NOMAD) are derived from (*53*) and are publicly available at https://doi.org/10.12770/3ea321a1-d9d4-49e5-a592-605b80dec240. Past and future model outputs are available upon request. Model-derived past and future front datasets are available upon request and will be made publicly available on Zenodo upon publication.

## Notes

https://github.com/IsaakBM/mmw_belmont

### Competing Interest Statement

The authors have declared no competing interest.

