## Supplementary Materials for "Megafauna show pervasive yet distinct affinity to ocean fronts: the urgent need for adaptive conservation in a warming world"

**This PDF file includes:**

Figs. S1 to S9
Table S1

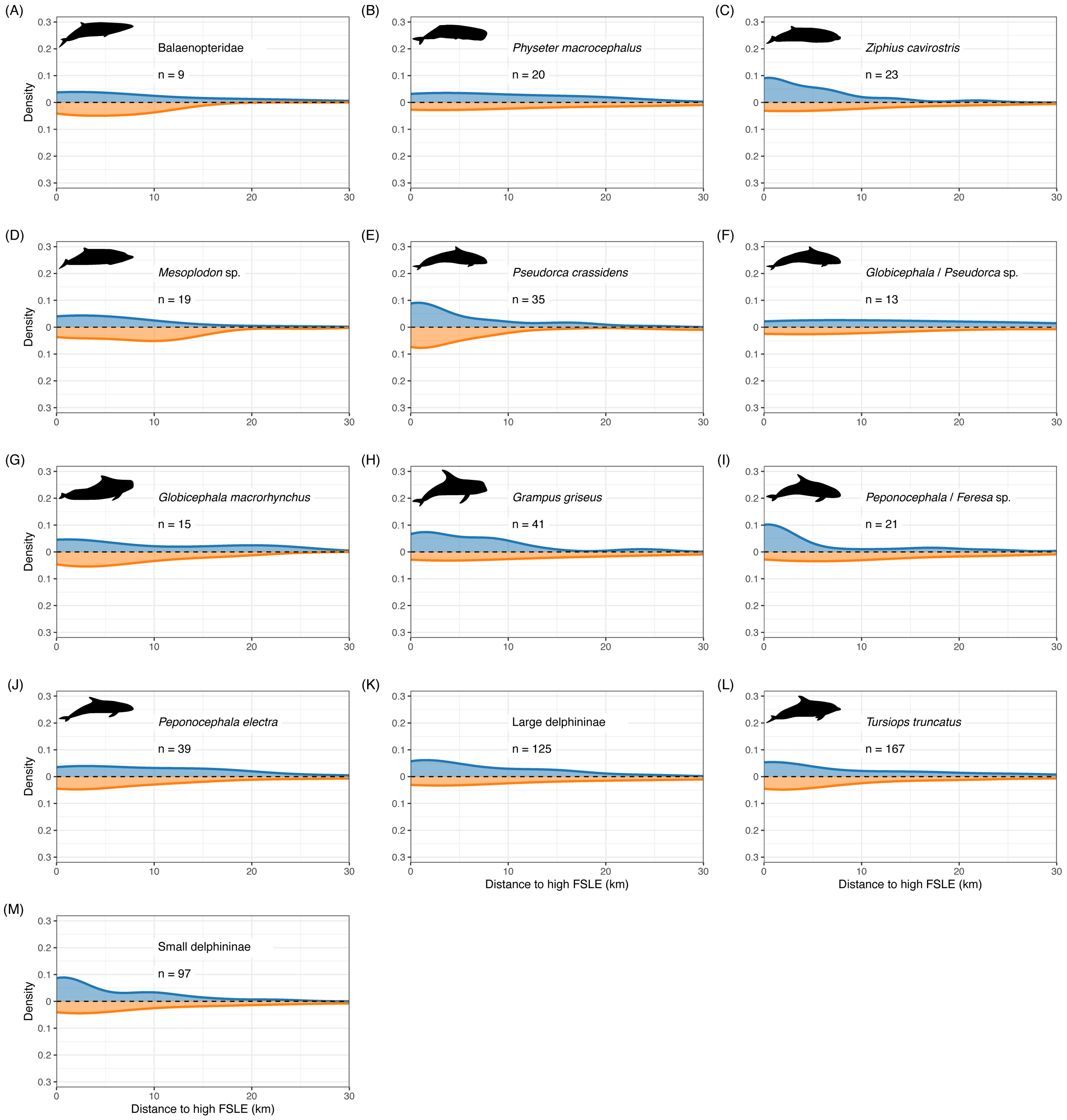

**Fig. S1.** Kernel density plots illustrate the relationship between density and distance to high (upper quartile) dynamical front areas for marine mammals. The blue distribution indicates the original data, and the orange distribution represents the distribution derived from the random point analysis. For each graph, the number of observations (n) has been included.

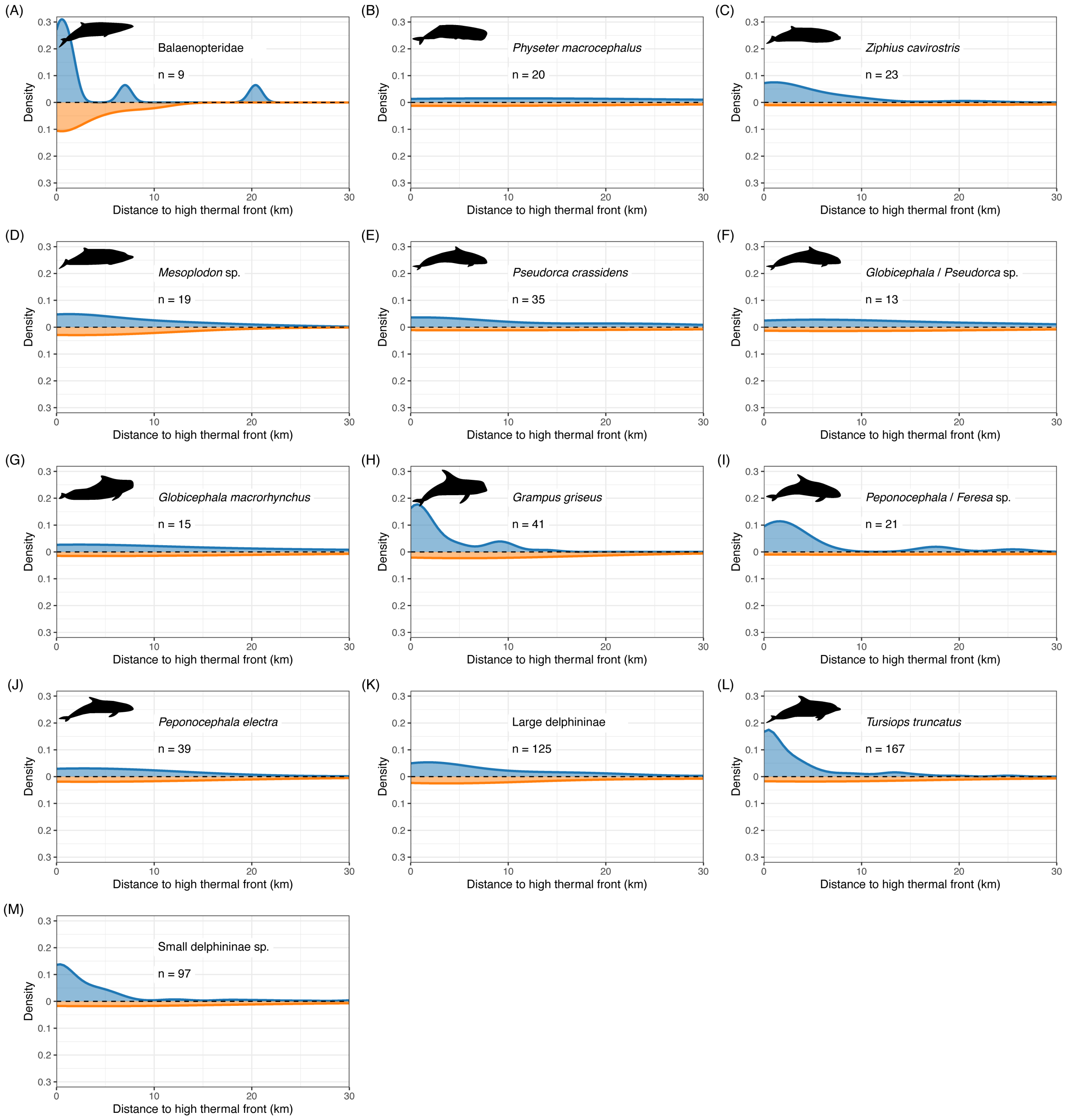

**Fig. S2.** Kernel density plots illustrate the relationship between density and distance to high (upper quartile) thermal front areas for marine mammals. The blue distribution indicates the original data, and the orange distribution represents the distribution derived from the random point analysis. For each graph, the number of observations (n) has been included.

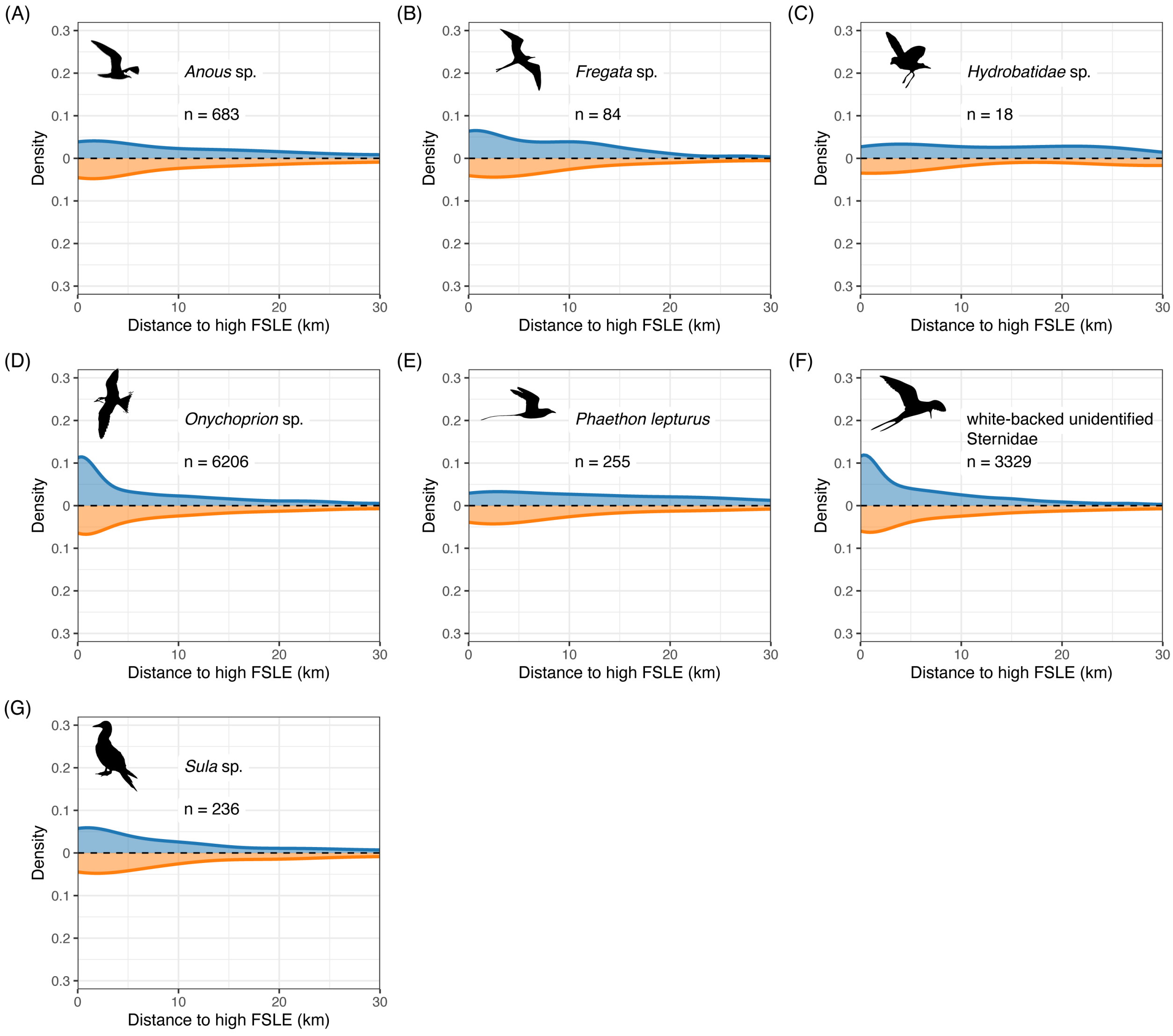

**Fig. S3.** Kernel density plots illustrate the relationship between density and distance to high (upper quartile) dynamical front areas for seabirds. The blue distribution indicates the original data, and the orange distribution represents the distribution derived from the random point analysis. For each graph, the number of observations (n) has been included.

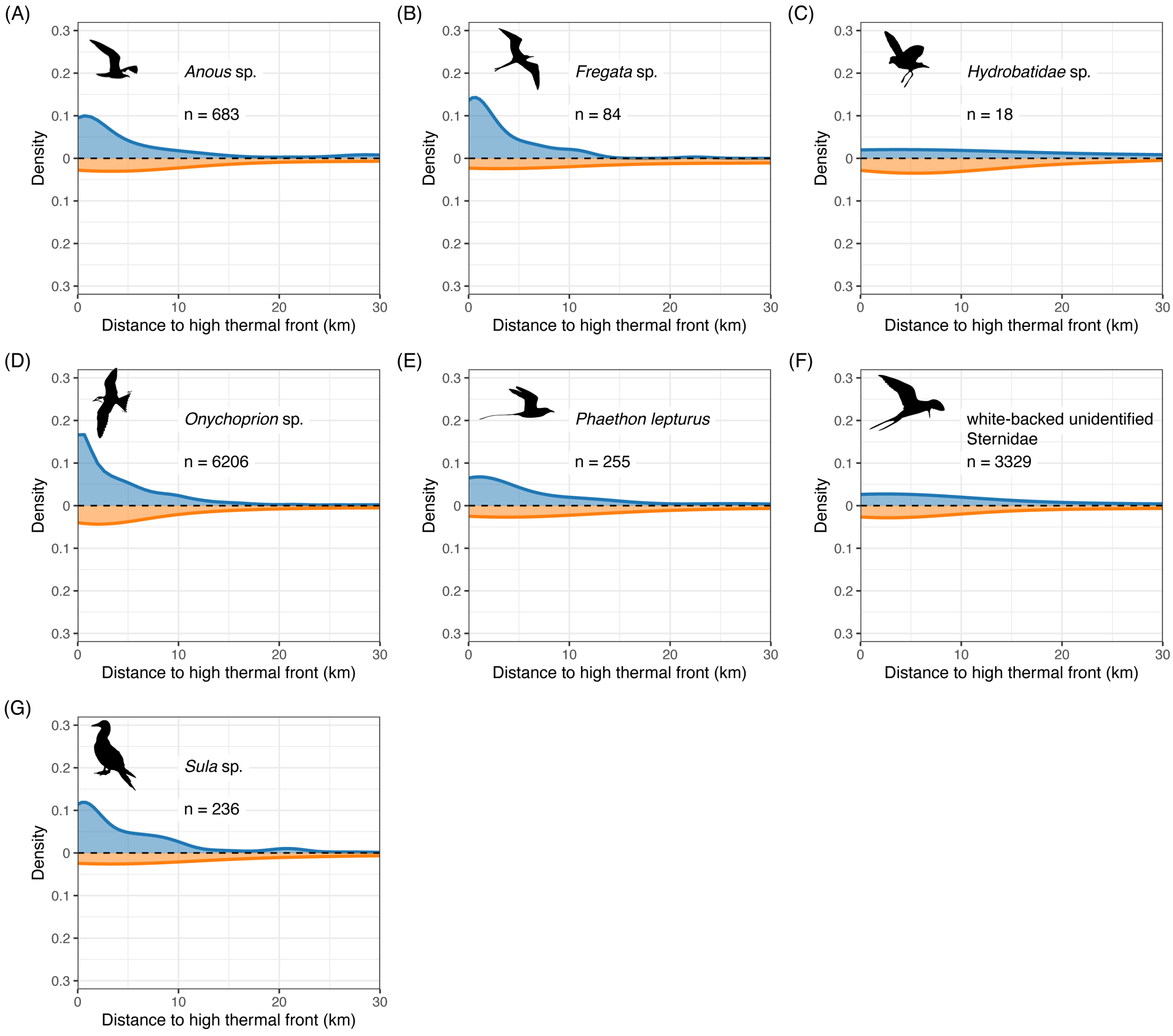

**Fig. S4.** Kernel density plots illustrate the relationship between density and distance to high (upper quartile) thermal front areas for seabirds. The blue distribution indicates the original data, and the orange distribution represents the distribution derived from the random point analysis. For each graph, the number of observations (n) has been included.

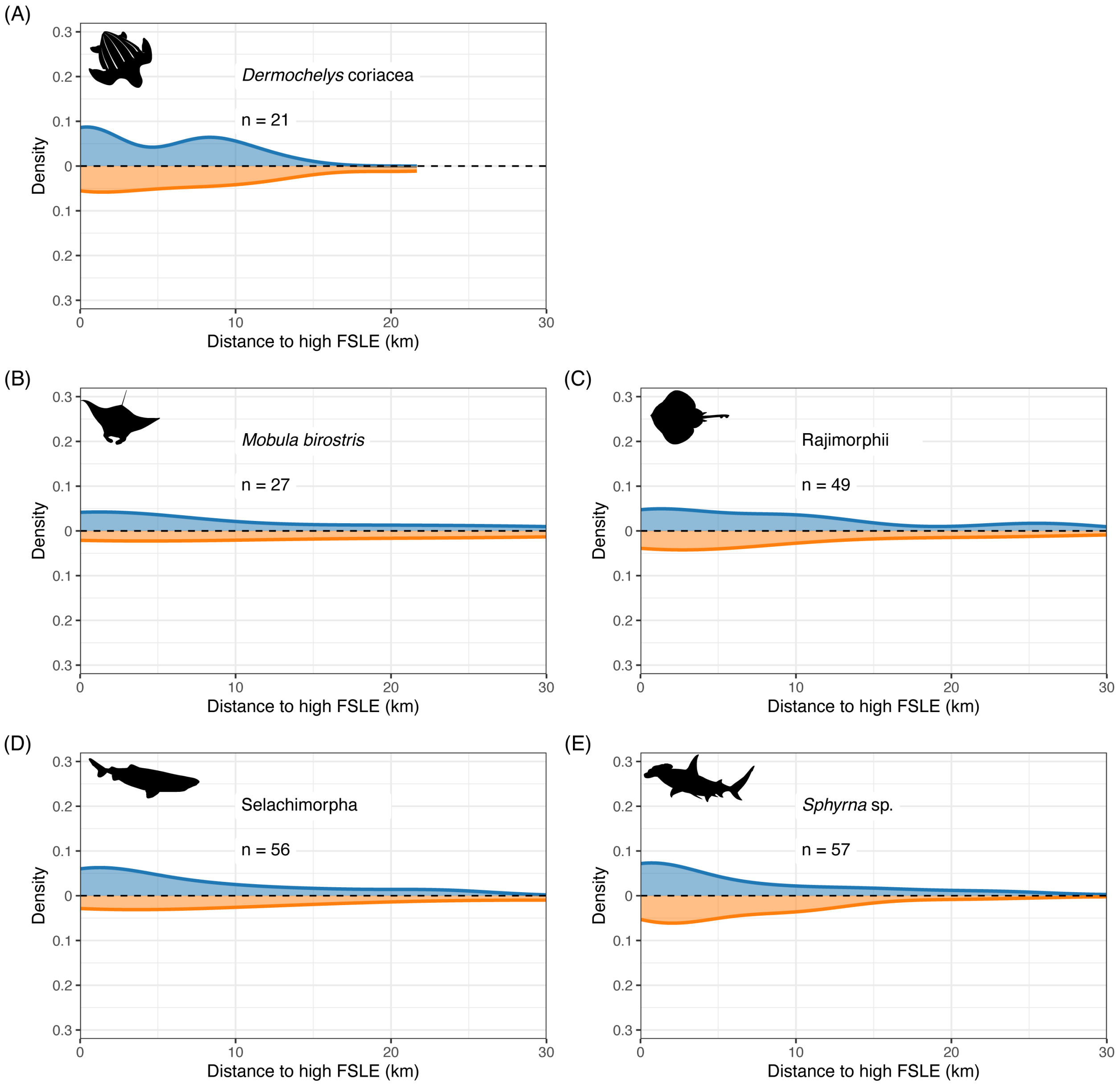

**Fig. S5.** Kernel density plots illustrate the relationship between density and distance to high (upper quartile) dynamical front areas for three different taxonomic groups: (A, B) hardturtles, (C, D) rays, and (E, F, G) sharks. The blue distribution indicates the original data, and the orange distribution represents the distribution derived from the random point analysis. For each graph, the number of observations (n) has been included.

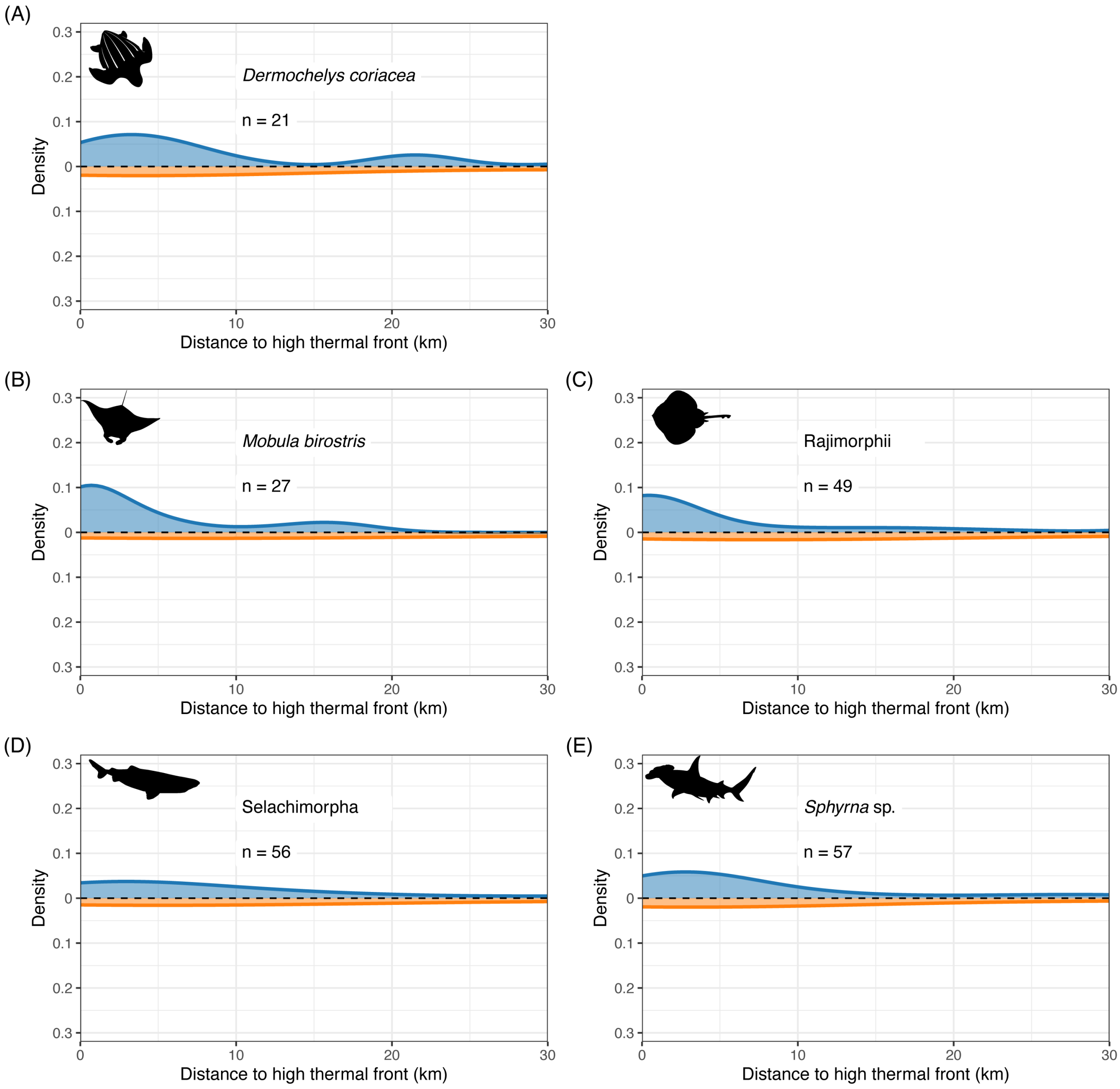

**Fig. S6.** Kernel density plots illustrate the relationship between density and distance to high (upper quartile) thermal front areas for three different taxonomic groups: (A, B) turtles, (C, D) rays, and (E, F, G) sharks. The blue distribution indicates the original data, and the orange distribution represents the distribution derived from the random point analysis. For each graph, the number of observations (n) has been included.

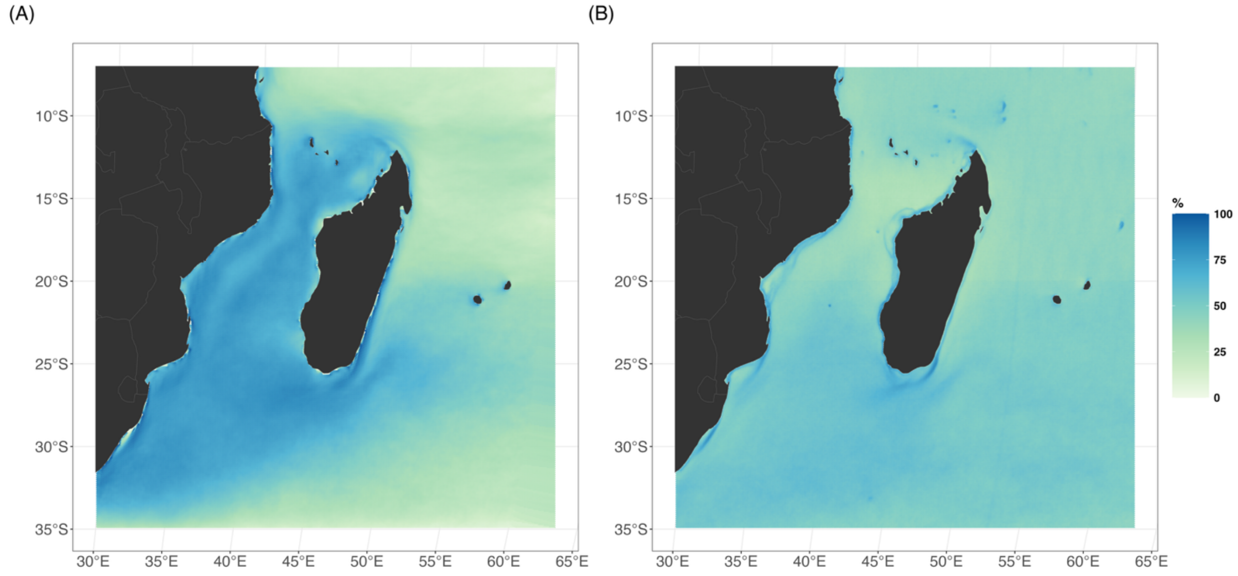

**Fig. S7.** Frequency of ocean fronts in the upper quartile derived from (A) Finite-Size Lyapunov Exponents (FSLE) from 1994 to 2021 and (B) Thermal Gradients from 2003 to 2021 across the South-Western Indian Ocean.

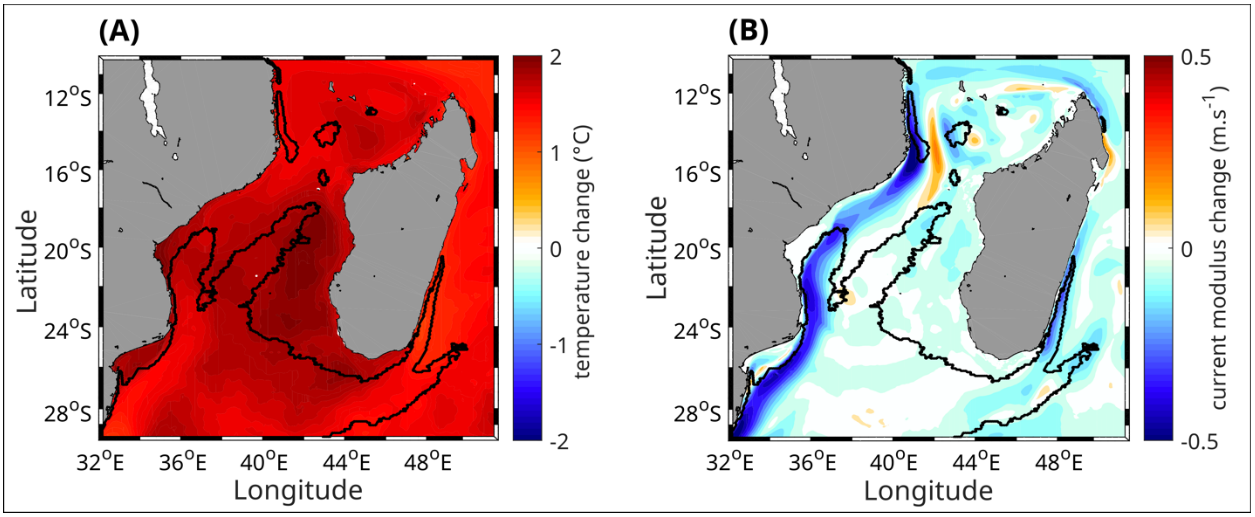

**Fig. S8.** Changes in mean sea surface temperature (A) and surface current speed (B) between hindcast (1993-2014) and pseudo global warming experiments (2076-2097). The bold black contour represents the area where both thermal and dynamical fronts are highly frequent in past decades (2003-2021), as shown in Fig. 3.

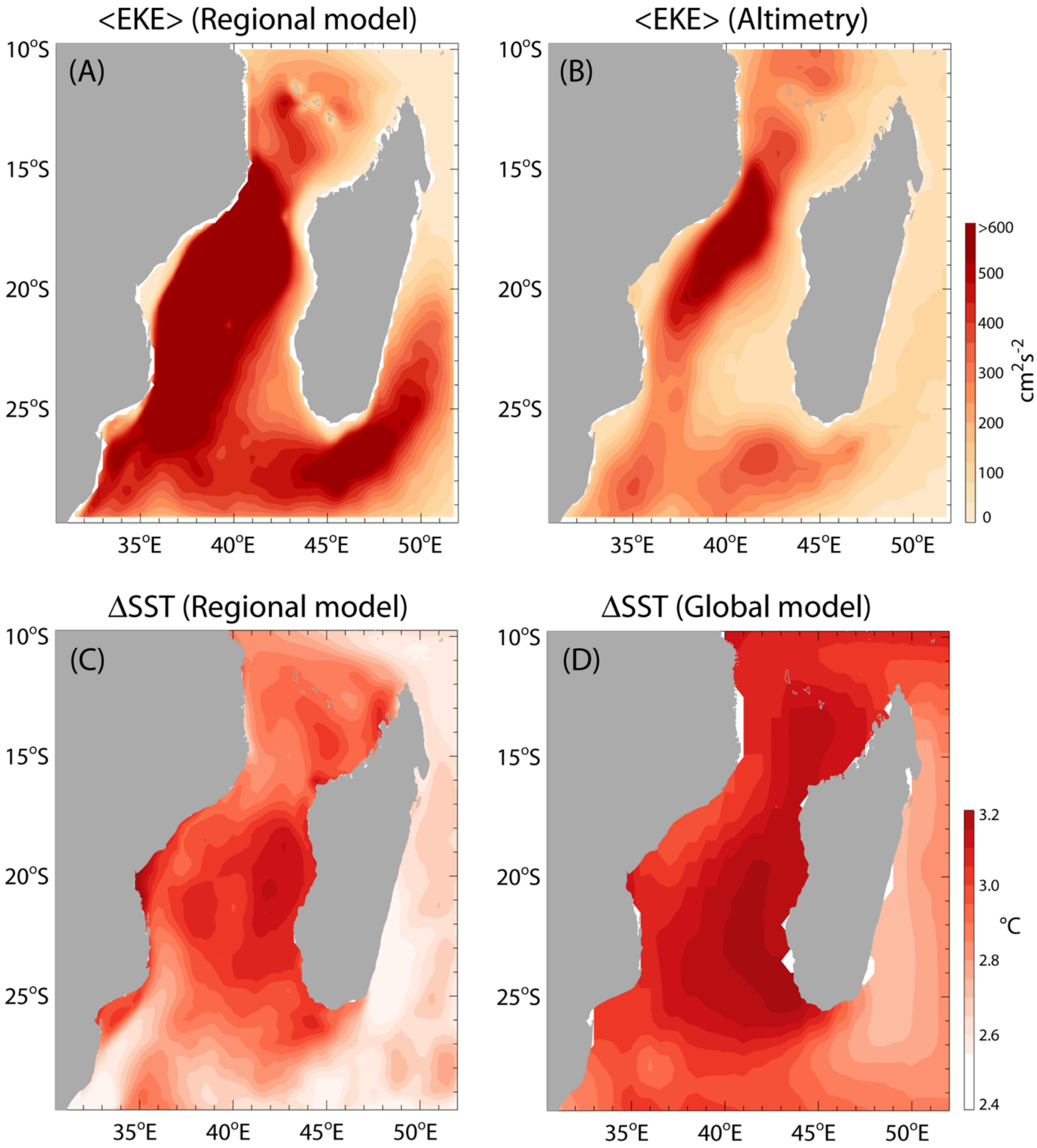

**Fig. S9. Model evaluation**: (top panels) mean eddy kinetic energy (in $cm^2s^{-2}$) computed from (A) the regional model control run and (B) the satellite-altimetry archive for the period 1993-2014. (bottom panels) Changes in Sea Surface Temperature (SST) between the future (2076-2097) and present (1993-2014) climates for the (C) regional and (D) global CESM-LENS models in °C.

**Table S1.** Taxonomic resolution and trophic regimes of marine megafauna species included in this study. Trophic regimes reflect general feeding habits such as plankton, small fish, squid, and larger prey like marine mammals or large fish, while trophic levels indicate the position of each species or group in the food chain. Values are derived from literature sources, and ranges indicate variability in diet composition or trophic interactions.

| Common/latin name | Taxonomic resolution | Trophic regime[1] | Trophic level[2] |
|---|---|---|---|
| **Tracking data** | | | |
| *Rhincodon typus* | Species | Plankton, small fish, squid | 3 |
| *Phaethon rubricauda* | Species | Small fish and squid | 3.5 – 4 |
| *Ardenna pacifica* | Species | Small fish and squid | 3.5 – 4 |
| *Caretta caretta* | Species | Omnivores | 2, but also 3 |
| **Marine mammals** | | | |
| *Balaenopteridae* | Family | Krill, small fishes, copepods | 3.5 |
| *Physeter macrocephalus* | Species | Large squid, large fish | 4.5 |
| *Ziphius cavirostris* | Species | | 4.5 |
| *Mesoplodon* sp. | Genus | | 4.4 |
| *Pseudorca crassidens* | Species | | 4.5 |
| *Globicephala* / *Pseudorca* sp. | Genus | | 4.5 |
| *Globicephala macrorhynchus* | Species | | |
| *Grampus griseus* | Species | | 4.1 – 4.3 |
| *Peponocephala* / *Feresa* sp. | Genus | Large squid, large fish | 4 – 4.5 |

| | | | |
|---|---|---|---|
| *Peponocephala electra* | Species | Large squid, large fish | 4 – 4.5 |
| Large delphininae | Family | | 4 – 4.5 |
| *Tursiops truncatus* | Species | | 4 – 4.5 |
| Small *delphininae* sp. | Family | | 4 – 4.5 |
| **Seabirds** | | | |
| *Anous* sp. | Genus | small fish, squid, and invertebrates | 3? |
| *Fregata* sp. | Genus | piscivorous | 4 (Ogden et al. 2014) |
| *Hydrobatidae* sp. | Genus | phytoplankton and small invertebrate | 3 |
| *Onychoprion* sp. | Genus | small fish, squid, and crustaceans | 3.5 – 4 |
| *Phaethon lepturus* | Species | Small fish and squid | 3.5 – 4 |
| *Phaethon* sp. | Genus | Small fish and squid | 3.5 – 4 |
| *Procellariidae* sp. | Genus | small fish, squid, and crustaceans | 3 – 4 |
| *Sternidae* sp. | Genus | small fish, squid, and crustaceans | 3 – 4 |
| *Sula* sp. | Genus | fish, squid | 4 |
| **Other Megafauna (turtles, rays, sharks)** | | | |
| *Cheloniidae* sp. | Genus | Omnivores | 2, but also 3 |
| *Dermochelys coriacea* | Species | jellyfish, salps, and soft-bodied invertebrates. | 3 – 4 |
| *Mobula* sp. | Species | | 3.5 |
| *Rajimorphii* sp. | Genus | crustaceans, mollusks, and small fish | 3.5 – 4 |
| *Rhincodon typus* | Species | Plankton, small fish, small squid | 3 |

| | | | |
|---|---|---|---|
| Selachimorpha | Superorder | fish, invertebrates, marine mammals and other sharks | 4 – 5 |
| *Sphyrna* sp. | Genus | fish, invertebrates, marine mammals and other sharks | 4 – 5 |

[1]Trophic regime: Herbivores, Planktivores, Detritivores, Omnivores, Carnivores, Piscivores, Apex predators
[2]Trophic level: 1 - 5
